# Reproducible big data science: A case study in continuous FAIRness

**DOI:** 10.1101/268755

**Authors:** Ravi Madduri, Kyle Chard, Mike D’ Arcy, Segun C. Jung, Alexis Rodriguez, Dinanath Sulakhe, Eric W. Deutsch, Cory Funk, Ben Heavner, Matthew Richards, Paul Shannon, Gustavo Glusman, Nathan Price, Carl Kesselman, Ian Foster

## Abstract

Big biomedical data create exciting opportunities for discovery, but make it difficult to capture analyses and outputs in forms that are findable, accessible, interoperable, and reusable (FAIR). In response, we describe tools that make it easy to capture, and assign identifiers to, data and code throughout the data lifecycle. We illustrate the use of these tools via a case study involving a multi-step analysis that creates an atlas of putative transcription factor binding sites from terabytes of ENCODE DNase I hypersensitive sites sequencing data. We show how the tools automate routine but complex tasks, capture analysis algorithms in understandable and reusable forms, and harness fast networks and powerful cloud computers to process data rapidly, all without sacrificing usability or reproducibility—thus ensuring that big data are not hard-to-(re)use data. We compare and contrast our approach with other approaches to big data analysis and reproducibility.

## 1 Introduction

Rapidly growing data collections create exciting opportunities for a new mode of scientific discovery in which alternative hypotheses are developed and tested against existing data, rather than by generating new data to validate a predetermined hypothesis [1,2]. A key enabler of these data-driven discovery methods is the ability to easily access and analyze data of unprecedented size, complexity, and generation rate (i.e., volume, variety, and velocity)—so called big data. Equally important to the scientific method is that results be easily consumed by other scientists [3, 4]: that is that results be findable, accessible, interoperable, and re-usable (FAIR) [5].

Yet there is currently a considerable gap between the scientific potential and practical realization of data-driven approaches in biomedical discovery [6]. This unfortunate situation results, in part at least, from inadequacies in the computational and data management approaches available to biomedical researchers. In particular, tools rarely scale to big data. For example, while a desktop tool may allow an analysis method to be readily applied to a small dataset (e.g., a single genome), applying the same method to a large dataset (e.g., 1,000 genomes) requires specialized infrastructure and expertise. The complexity of the associated data and computation management tasks frequently becomes a gating factor to progress. These difficulties are magnified by the disjointed nature of the biomedical data landscape, which often lacks common interfaces for data discovery and data access, conventions for bundling and transporting datasets, and methods for referencing data produced in different locations,. and features non-portable and idiosyncratic analysis suites.

We show here that these difficulties can be overcome via the use of relatively simple tools that either entirely automate or significantly streamline the many, often mundane, tasks that consume biomedical researcher time. These tools include Big Data Bags (BDBags) for data exchange and minimal identifiers (Minids) as persistent identifiers for intermediate data products [7]; Globus cloud services for authentication and data transfer [8,9]; and the Galaxy-based Globus Genomics [10] and Docker containers [11] for reproducible cloud-based computations. Simple application programming interface (API)-level integration means that, for example, whenever a new BDBag is created to bundle outputs from a computation, a Minid can easily be created that can then be consumed by a subsequent computational step.

To demonstrate what can be achieved in this space, we present here a case study of big data analysis, a transcription factor binding site (TFBS) analysis that creates an atlas of putative transcription factor binding sites from ENCODE DNase I hypersensitive sites sequencing (DNase-seq) data, across 27 tissue types. This application involves the retrieval and analysis of multiple terabytes of publicly available DNase-seq data with an aggregated set of position weight matrices representing transcription factor binding sites; a range of open source analysis programs, Galaxy workflows, and customized R scripts; high-speed networks for data exchange; and tens of thousands of core-hours of computation on workstations and public clouds. We introduce the analysis method, review the tools used in its implementation, and present the implementation itself, showing how the tools enable the principled capture of a complex computational workflow in a reusable form. In particular, we show how all resources used in this work, and the end-to-end process itself, are captured in reusable forms that are accessible via persistent identifiers.

The remainder of this paper is as follows. In §2, we introduce the TFBS atlas application and in §3 the tools that we use to create a FAIR implementation. We describe the implementation in §4 and §5, discuss implications of this work and its relationship to other approaches in §6, and conclude in §7.

## 2 An atlas of transcription factor binding sites

Large quantities of DNase I hypersensitive sites sequencing (DNase-seq) data are now available, for example from the Encyclopedia of DNA Elements (ENCODE) [12]. Funk et al. [13] show how such data can be used to construct genome-wide maps of candidate transcription factor binding sites (TFBSs) via the large-scale application of footprinting methods. As outlined in Fig 1, their method comprises five main steps, which are labeled in the figure and referenced throughout this paper as ❶‥❺:

❶ Retrieve tissue-specific DNase-seq data from ENCODE, for hundreds of biosample replicates and 27 tissue types.
❷ Combine the DNase-seq replicates data for each aligned replicate in each tissue and merge the results. Alignments are computed for two seed sizes, yielding double the number of output files.
❸ Apply two footprinting methods—Wellington [14] and HMM-based identification of TF footprints (HINT) [15], each of which has distinct strengths and limitations [16]—to each DNase-seq from ❷ to infer footprints. (On average, this process identifies a few million footprints for each tissue type, of which many but certainly not all are found by both approaches.)
❹ Starting with a supplied set of non-redundant position weight matrices (PWMs) representing transcription-factor-DNA interactions, create a catalog of “hits” within the human genome, i.e., the genomic coordinates of occurrences of the supplied PWMs.
❺ Intersect the footprints from ❸ and the hits from ❹ to identify candidate TFBSs in the DNase-seq data.

**Fig 1.**
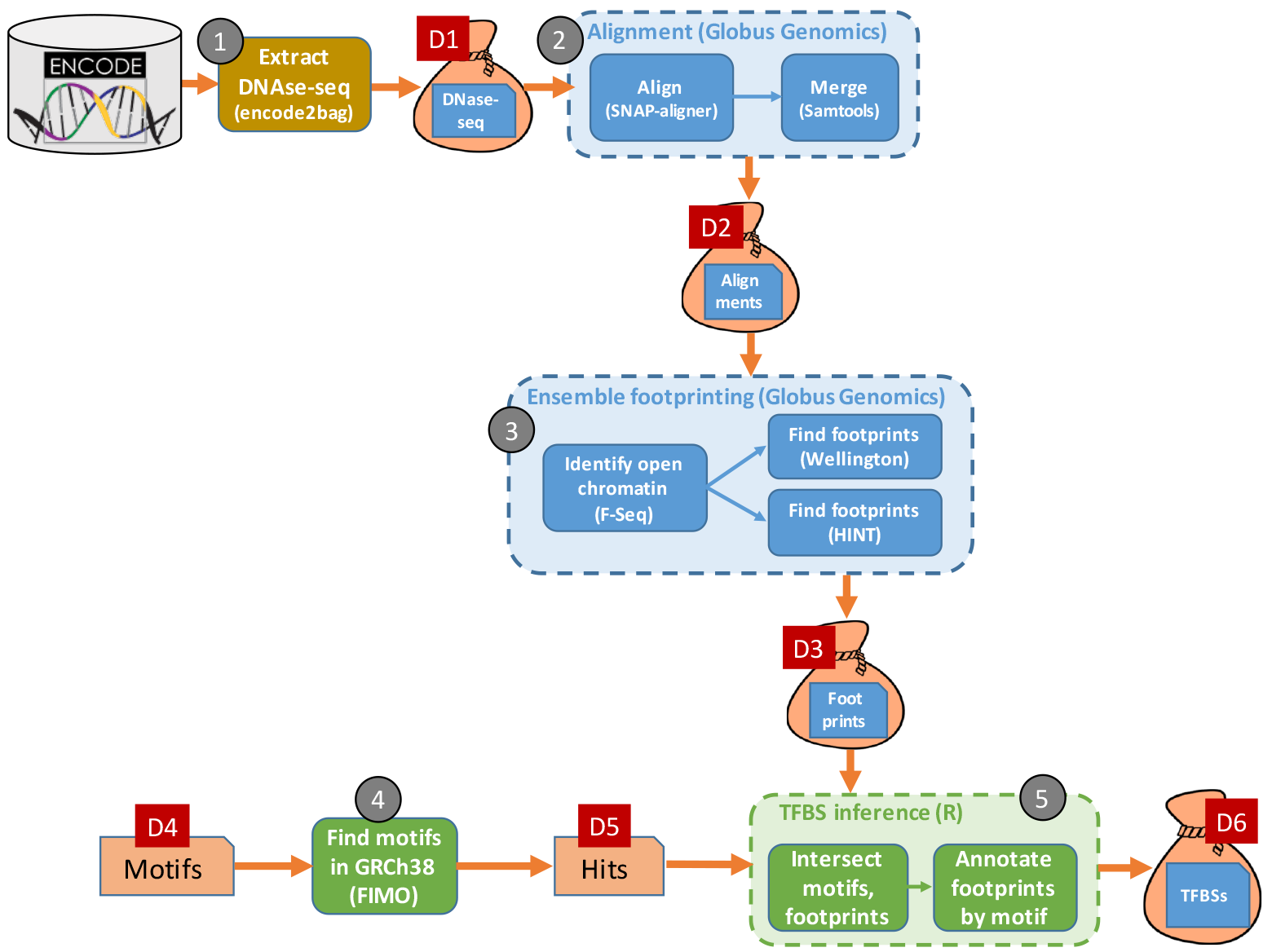
A high-level view of the TFBS identification workflow, showing the six principal datasets, labeled D1-D6, and the five computational phases, labeled ❶–❺

We provide more details on each step in subsequent sections, where we also discuss the specifics of the data that are passed between steps and preserved for subsequent access. Here we note some characteristics of the overall workflow that are important from a reproducibility perspective. The workflow involves many steps and files, making it important that the provenance of final results be captured automatically rather than manually. It involves considerable data (terabytes: TBs) and computation (hundreds of node-hours on 32-core nodes) and thus requires parallel computing (e.g., on a cloud) in order to complete in a timely manner. Finally, it makes use of multiple types of computation: an online service (**`encode2bag`**), big data Galaxy pipelines running in parallel on the cloud, and R code running on a workstation or laptop. These diverse characteristics are typical of many modern bioinformatics applications.

### 3 Tools used in TFBS atlas implementation

Before describing our implementation of the TFBS workflow, we introduce tools that we leverage in its development. These tools, developed or enhanced within the NIH-funded Big Data for Discovery Science center (BDDS) [17], simplify the development of scalable and reusable software by providing robust solutions to a range of big data problems, from data exchange to scalable analysis.

#### 3.1 BDBags, Research Objects, and Minids for data exchange

Reproducible big data science requires mechanisms for describing, referring to, and exchanging large and complex datasets that may comprise many directories and files (elements). Key requirements here are [7] enumeration: explicit enumeration of a dataset’s elements, so that subsequent additions or deletions can be detected; fixity: enable verification of dataset contents, so that data consumers can detect errors in data transmission or modifications to data elements; description: provide interoperable methods for tracking the attributes (metadata) and origins (provenance) of dataset contents; distribution: allow a dataset to contain elements from more than one location; and identification: provide a reliable and concise way of referring to datasets for purposes of collaboration, publication, and citation.

We leverage three technologies to meet these requirements. We use the BDBag to define a dataset and its contents by enumerating its elements, regardless of their location (enumeration, fixity, and distribution); the Research Object (RO) [18] to characterize a dataset and its contents with arbitrary levels of detail, regardless of their location (description); and the Minid to uniquely identify a dataset and, if desired, its constituent elements, regardless of their location (identify, fixity). Importantly, these mechanisms can all be used without deploying complex software on researcher computers.

The **Big Data Bag** (BDBag) exchange format and associated tools [7] provide a robust mechanism for exchanging large and complex data collections. The BDBag exchange format extends the BagIt specification [19] to provide a self-describing format for representing large data. To give a flavor of the BDBag specification, we show an example in Fig 2. The dataset contents are the directories and files contained within the data directory. The other elements provide checksum and metadata information required to verify and interpret the data. Importantly, a BDBag can encode references to remotely accessible data, with the information required to retrieve those data provided in the fetch.txt file as a (URL, LENGTH, FILENAME) triple. This mechanisms supports the exchange of large datasets without copying large amounts of data (a “holey” BDBag that contains only references, not data, may be just a few hundreds of bytes in size); it also allows the definition of data collections that specific individuals may not have immediate permission to access, as when access to data elements is restricted by data use agreements. Given a BDBag, BDBag tools can be used to materialize those data in a standard way, capture metadata, and verify contents based on checksums of individual data and metadata items.

**Fig 2.**
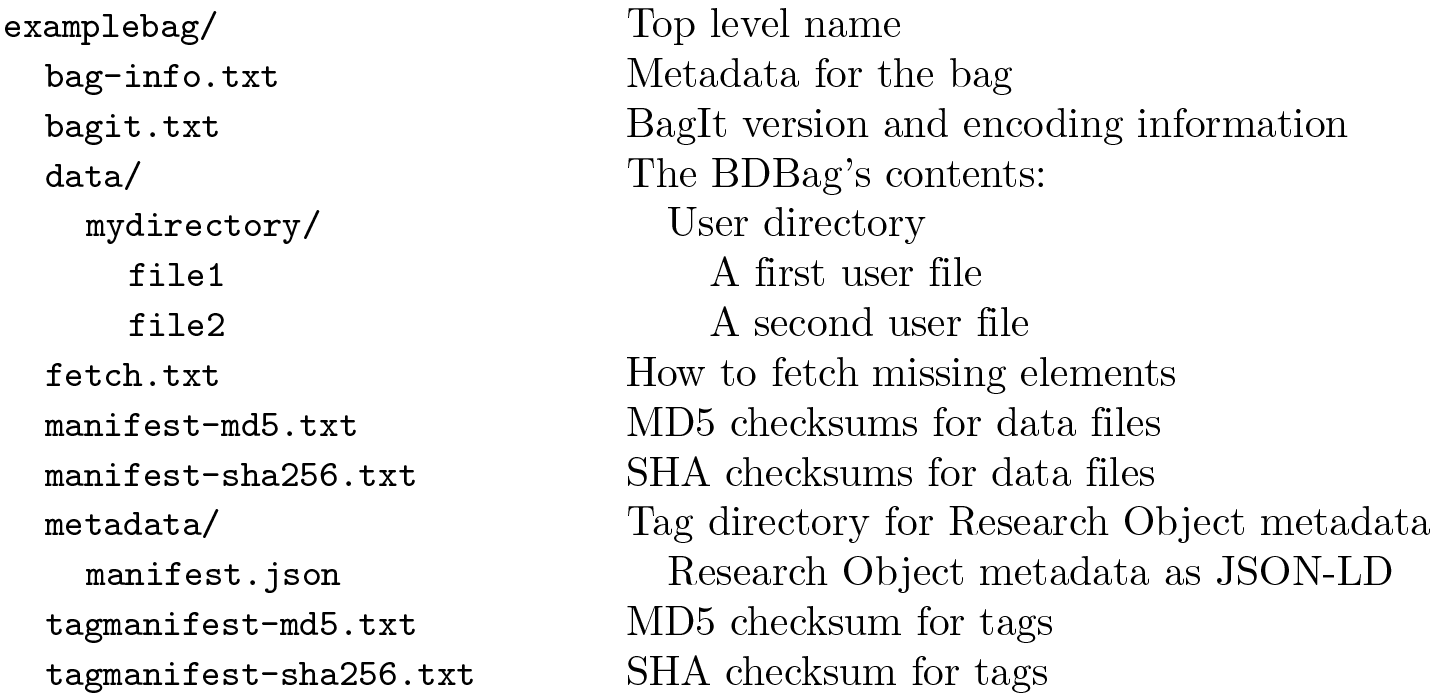
An example BDBag, with contents in the **`data`** folder, description in the **`metadata`** folder, and other elements providing data required to fetch remote elements (**`fetch.txt`**) and validate its components.

The BDBag specification adopts the **Research Object** (RO) framework to associate attribution and provenance information, and structured and unstructured annotations describing individual resources, with the data contained in a BDBag. A BDBag’s metadata directory contains annotations and the RO manifest.json in JSON-LD format [20]: see Fig 2.

Reproducible science requires mechanisms for robustly naming datasets, so that researchers can uniquely reference and locate data, and share and exchange names (rather than an entire dataset) while being able to ensure that a dataset’s contents are unchanged. We use the minimal viable identifier (**Minid**) [7] for this purpose. As the name suggests, Minids are lightweight identifiers that can be easily created, resolved, and used. Minids take the form **`minid:[suffix]`**, where the suffix is a unique sequence of characters. The **minid** prefix is registered at identifiers.org and n2t.net, so that, for example, visiting the URL https://n2t.net/minid:b9q119) redirects the browser to the landing page for the object with identifier **`minid:b9q119:`** see Fig 3.

**Fig 3.**
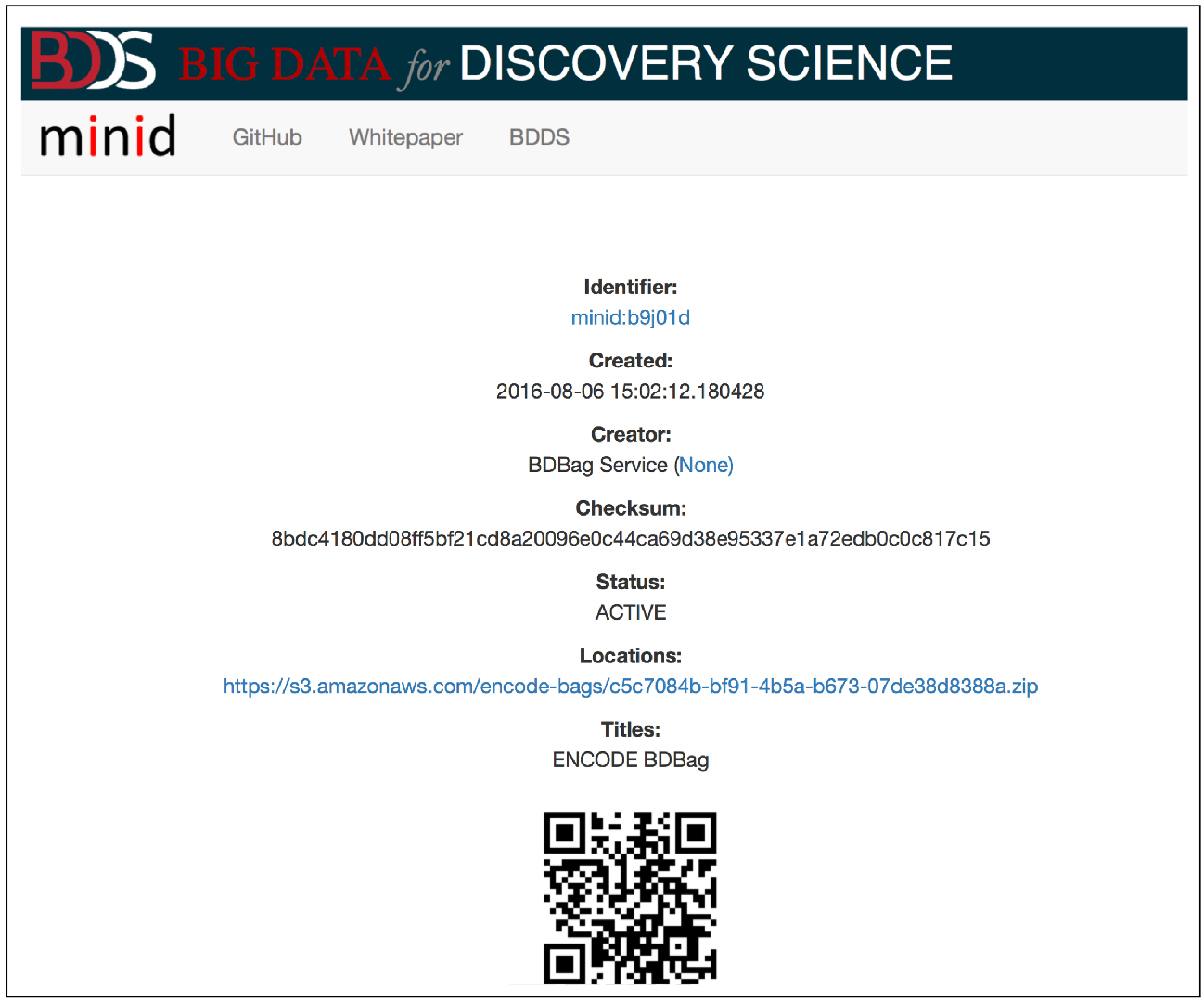
A Minid landing page for a BDBag generated by the **`encode2bag`** tool, showing the associated metadata, including locations (in this case, just one).

A landing page is intended to be always available, even if the data are not. It presents the complete metadata record for the Minid and links to one or more locations where the referenced data can be obtained. Allowing more than one data location enables data replication. For example, a Minid may reference one copy of a dataset in the source repository, additional copies in different vendor cloud object stores, and yet another copy on a local file system. Because the Minid includes a checksum of the content, we can ensure that whatever copy is used, it contains the correct and intended content. It is up to the consumer of the Minid to determine which copy of the data is “best” for their purpose, and the responsibility of the Minid creator to interact with the landing page service to register new copies. The GET methods for the landing page support HTTP content negotiation; results may be returned in human-readable (HTML) or machine-readable (JSON) form.

While Minids and BDBags can be used independently, they can be used together to powerful effect. As we illustrate in later sections, we can create a Minid for a BDBag, allowing us to uniquely identify the BDBag instance and providing a repeatable method for referring to the BDBag. A Minid can be used as the URL for a remote file reference within a BDBag’s **`fetch.txt`** file, in place of a direct URL to a file storage location. The BDBag tooling knows how to resolve such a Minid reference through the landing page to a copy of the BDBag data for materialization into the complete data set. We leverage both of these combinations of Minids and bags in the TFBS workflows.

#### 3.2 Globus data management services

The often distributed nature and large size of biomedical data complicates data management tasks—such as, in our case study, moving ENCODE data to cloud computers for analysis, and providing persistent access to analysis results. We use two capabilities provided by Globus [9] to overcome these difficulties.

First, we use Globus identity management, authentication, and authorization capabilities to enable researchers to authenticate with their institutional credentials and then access different data sources and data services without requiring further authentication.

Second, we use the Globus file-based data management services to enable efficient, reliable, and secure remote data access, secure and reliable file transfer, and controlled sharing. With more than 10,000 storage systems accessible via the Globus Connect interface, and support for data access from many different file systems and object stores, Globus translates the often baffling and heterogeneous world of distributed storage into a uniform, easily navigable data space.

A third capability that we expect to leverage in future work is Globus data publication [21] to support large-scale data publication. This service provides workflows for making data immutable, associating descriptive metadata, and assigning persistent identifiers such as digital object identifiers (DOIs) [22].

#### 3.3 Globus Genomics for parallel cloud-based computation

Small data analyses can be implemented effectively via R or Python scripts, that can be executed on a workstation or a cloud-hosted virtual machine and then shared as documents or via notebook environments such as Jupyter [23]. Big data analyses can be more challenging to implement and share, due to the need to orchestrate the execution of multiple application programs on many processors in order to process large quantities of data in a timely manner, whether for quality control [24], computation of derived quantities, or other purposes.

We use Globus Genomics [10] to orchestrate multi-application analysis on multi-processor cloud computing platforms. Globus Genomics builds on the Galaxy workflow environment [25], widely used in bioinformatics to support the graphical specification of multi-step computational analyses. Globus Genomics extends Galaxy’s capabilities with support for Globus data access, parallel execution on cloud platforms, dispatch of applications within Docker containers, input of BDBags referenced via Minids, and other features useful for big data applications.

Other workflow systems with capabilities similar to those of the Galaxy system include the Python-based Toil [26], the Pipeline environment [27,28], and the Common Workflow Language (CWL) [29]. The approach described here could be easily adopted to use different workflow languages and systems.

#### 3.4 Containers for capturing software used in computations

A final challenge in reproducible science is recording the software used to perform a computation. Source code allows a reader to examine application logic [30, 31], but may not run on a new platform. Container technologies such as Docker [11] and Singularity [32] can be used to capture a complete software stack in a form that can be executed on many platforms. We use Docker here to package the various applications used in the footprinting workflow. A benefit of this technology is that a container image can be described (and built) with a simple text script that describes the base operating system and the components to be loaded: in the case of Docker, a *Dockerfile*. Thus it is straightforward to version, share, and reproducibly rebuild a container [33].

#### 4 A scalable, reproducible TFBS workflow

Having described the major technologies on which we build, we now describe the end-to-end workflow of Fig 1. We cover each of ❶ – ❺ in turn. Table 1 summarizes the biosamples, data, and computations involved in the workflow.

**Table 1.** Details of the per-tissue computations performed in the ensemble footprinting phase. Data sizes are in GB. Times are in hours on a 32-core AWS node; they sum to 2,149.1 node hours or 68,771 core hours. DNase: DNase Hypersensitivity (DNase-seq) data from ENCODE. Align: Aligned sequence data. Foot: Footprint data and footprint inference computation. Numbers may not sum perfectly due to rounding.

| Tissue | Biosamples | Replicates | Data size |  |  | Compute time |  |
| --- | --- | --- | --- | --- | --- | --- | --- |
|  |  |  | DNase | Align | Foot | Align | Foot |
| adrenal gland | 3 | 8 | 31 | 68 | 0.5 | 7.4 | 18.7 |
| blood vessel | 10 | 129 | 117 | 234 | 2.1 | 32.7 | 68.9 |
| bone element | 1 | 7 | 4 | 6 | 0.2 | 1.4 | 3.1 |
| brain | 29 | 185 | 402 | 840 | 6.2 | 120.0 | 160.5 |
| bronchus | 2 | 9 | 18 | 36 | 0.4 | 3.1 | 9.7 |
| esophagus | 2 | 41 | 35 | 64 | 0.3 | 12.6 | 7.7 |
| extraembryonic | 11 | 66 | 193 | 412 | 3.0 | 46.5 | 89.9 |
| eye | 8 | 53 | 129 | 252 | 2.3 | 26.9 | 109.5 |
| gonad | 2 | 7 | 27 | 56 | 0.4 | 4.9 | 12.2 |
| heart | 8 | 69 | 169 | 342 | 2.0 | 38.8 | 76.0 |
| kidney | 8 | 29 | 84 | 174 | 2.0 | 19.5 | 96.5 |
| large intestine | 5 | 18 | 96 | 184 | 0.9 | 23.2 | 50.0 |
| liver | 3 | 8 | 26 | 62 | 0.5 | 4.9 | 19.8 |
| lung | 7 | 94 | 142 | 300 | 1.8 | 19.3 | 27.2 |
| lymphatic vessel | 2 | 30 | 21 | 286 | 0.4 | 3.6 | 11.9 |
| lymphoblast | 21 | 71 | 150 | 320 | 2.9 | 70.0 | 153.5 |
| mammary gland | 2 | 5 | 18 | 36 | 0.4 | 2.8 | 10.7 |
| mouth | 4 | 18 | 81 | 164 | 1.1 | 19.5 | 57.3 |
| muscle organ | 4 | 13 | 54 | 110 | 0.8 | 9.6 | 49.6 |
| pancreas | 2 | 13 | 38 | 84 | 0.5 | 8.1 | 30.9 |
| prostate gland | 2 | 8 | 23 | 50 | 0.3 | 4.2 | 78.1 |
| skin | 48 | 401 | 377 | 780 | 8.8 | 220.0 | 181.2 |
| spinal cord | 2 | 34 | 66 | 128 | 0.7 | 9.2 | 28.0 |
| stomach | 1 | 5 | 24 | 52 | 0.4 | 3.5 | 3.6 |
| thyroid gland | 3 | 24 | 63 | 136 | 0.7 | 13.6 | 27.0 |
| tongue | 2 | 8 | 51 | 106 | 0.7 | 13.7 | 26.1 |
| urinary bladder | 1 | 2 | 4 | 8 | 0.2 | 0.8 | 1.8 |
| **Total** | **193** | **1,355** | **2,443** | **5,291** | **40.8** | **739.7** | **1,409.4** |

##### 4.1 Obtaining input data: encode2bag

The TFBS algorithm operates on DNase Hypersensitivity (DHS) data in the form of DNase-seq data obtained by querying the ENCODE database for DNase-seq data for each of 27 tissue types. These queries, when performed by Funk et al. [13], yielded a total of 1,591 FASTQ files corresponding to 1,355 replicates from 193 biosamples. (Each tissue-specific biosample may have multiple replicates: for example, the tissue type *adrenal gland* has eight replicates from three biosamples. Also, some replicates are broken into more than one FASTQ files.) Note that an identical query performed against ENCODE at a different time may yield different results, as new data are added or removed and quality control procedures evolve. Thus, it is important to record the results of these queries at the time they were executed, in a reproducible form.

ENCODE provides a web portal that a researcher can use to query the ENCODE database, using menus to specify parameters such as assay, biosample, and genomic annotations. The result is a set of data URLs, which must be downloaded individually and unfortunately do not come with associated metadata or context. Researchers often resort to building shell scripts to download and store the raw datasets. These manual data retrieval and management steps can be error-prone, time consuming, and difficult to reproduce. Researchers must manually save queries to record data provenance, and the only way to validate that downloaded files have not been corrupted is to download them again.

To simplify this process, we used BDBag, Minid, and Globus tools to create a lightweight command line utility and web service, **`encode2bag`**, shown as ❶ in Fig 1. A researcher can use either the web interface or the command line interface to either enter an ENCODE query or upload an ENCODE metadata file describing a collection of datasets. They can then access the corresponding data, *plus associated metadata and checksums*, as a BDBag. Fig 4 shows an illustrative example in which the web interface is used to request data from urinary bladder DNase-seq experiments.

**Fig 4.**
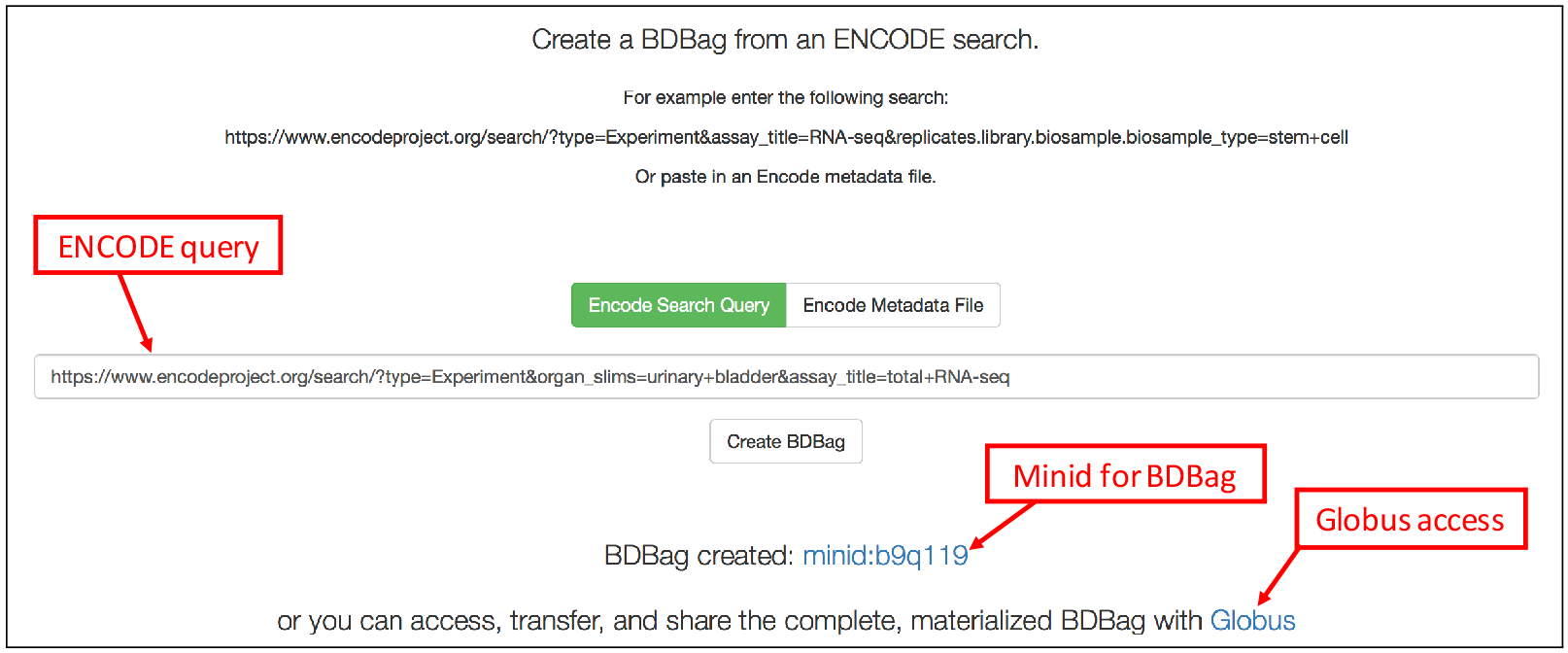
The **`ncode2bag`** portal. The user has entered an ENCODE query for urinary bladder DNase-seq data and clicked “Create BDBag.” The portal generates a Minid for the BDBag and a Globus link for reliable, high-speed access.

Selecting the “Create BDBag” button triggers the creation of a ~100 kilobyte BDBag that encapsulates references to the files in question, metadata associated with those files, the query used to identify the data, and the checksums required to validate the files and metadata. The BDBag is stored in AWS Simple Storage Service (S3) cloud storage from where it can be accessed for purposes of sharing, reproducibility, or validation. Because this BDBag contains references to data, rather the data themselves, it captures the entire response to the query in a small (hundreds of kilobytes) form that can be easily downloaded, moved, and shared. When needed, all, or a subset of, the files named within the BDBag’s fetch.txt file can be downloaded (using BDBag tools), while ensuring that their contents match those of the original query.

To further streamline access to query results, **`encode2bag`** assigns a Minid for each BDBag that it creates, so as to provide for unambiguous naming and identification of research data products that are used for data provenance. In the example in Fig 4 the Minid is **`minid:b9ql19`**; as discussed earlier, resolving this identifier leads to a landing page similar to that shown in Fig 3, which in turn contains a reference to the BDBag. The Minid can be passed between services as a concise reference to the BDBag.

The Funk et al. [13] workflow uses **`encode2bag`** to create BDBags for each of the 27 tissue types in ENCODE, each with its own Minid. For example, the DNase-seq data associated with adrenal tissue is at **`minid:b9w37t`**. These 27 BDBags contain references to a total of 2.4 TB of ENCODE data; references that can be followed at any time to access the associated data. It is these BDBags that are the input to the next step of the TFBS workflow,❷.

The fact that ❶ produces one BDBag per tissue type, each with a Minid, allows each tissue type to be processed independently in subsequent steps, providing considerable opportunities for acceleration via parallel processing. When publishing data for use by others, on the other hand, it would be cumbersome to share 27 separate Minids. Thus, as described in later sections, we also create for each such collection of BDBags a “bag of bags,” a BDBag that contains references to a set of other BDBags. This pervasive use of Minids and BDBags greatly simplifies the implementation and documentation of the TFBS workflow.

##### 4.2 Aligning DNase-seq sample data

Now that ❶ has prepared the input data, ❷ prepares those data for the footprinting computation. For each of the 27 tissue types, the input to this phase comprises DNase-seq replicates for one or more biosamples (typically multiple replicates for each biosample), organized as a BDBag. The analysis first fetches the sequence data and, for each biosample, uses the SNAP sequence aligner to align the associated replicates. The resulting replicate alignments are merged and sorted to yield a single binary alignment data (BAM) file per biosample. The BAM files for all biosamples for each tissue type are combined into a BDBag.

As the ENCODE data consist primarily of short sequence reads, Funk et al. [13] ran the sequence alignment process twice, with seed lengths of 16 and 20, respectively. ❶ thus produces two BDBags per tissue type, for a total of 54. (The two sets of outputs allow ❺ to compare the merits of the two seed lengths for identifying footprints.)

While the computations involved in ❷ are relatively simple, the size of the datasets being manipulated and the cost of the computations make it important to execute subcomputations in parallel whenever possible. Each tissue and seed can be processed independently, as can the alignments of the replicates for each biosample, the merge and sort for each biosample, and (in ❸) the footprint generation by HINT and Wellington. We use Globus Genomics to manage the resulting parallel computations in a way that both enables cloud-based parallel execution and reproducibility.

Fig 5 shows the Galaxy workflow that implements ❷ and ❸. In this figure, each smaller box represents a separate application, with a name (the shaded header), one or more inputs (below the header and above the line), and one or more outputs (below the line). Each link connects an output from one application to the input of another.

**Fig 5.**
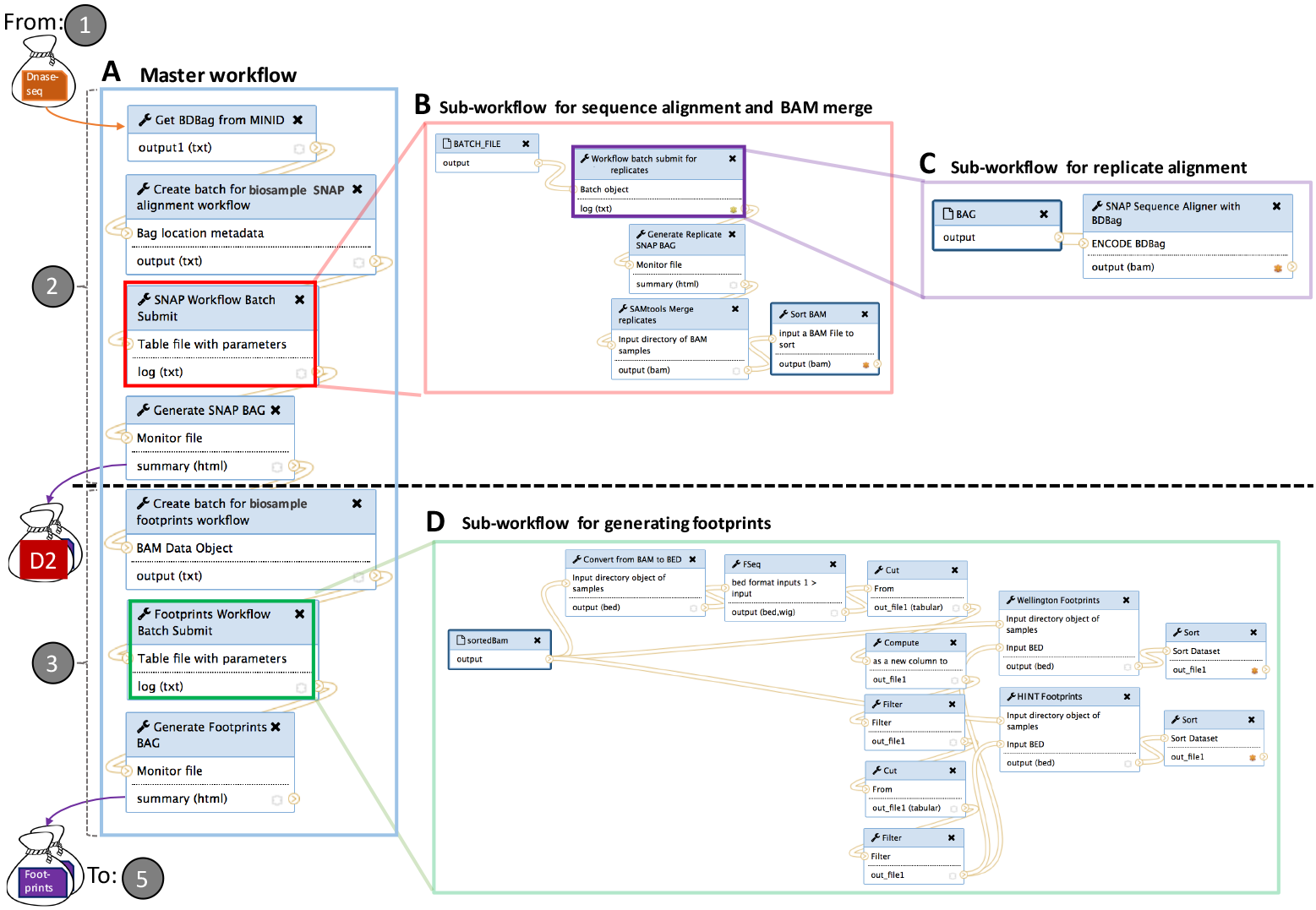
Our DNase-seq ensemble footprinting workflow, used to implement ❷ and ❸ of Fig 1. The master workflow A takes a BDBag from ❶ as input. It executes from top to bottom, using subworkflows B and C to implement ❷ and then subworkflow D to implement ❸. It produces as output BDBags containing aligned DNase-seq data and footprints, with the latter serving as input to ❺.

Fig 5 comprises four distinct workflows. The (A) master workflow, on the left, is run once for each tissue type, with each run taking a BDBag from ❶ as input and producing multiple BDBags as output. This workflow runs seven different applications in sequence, from top to bottom. Two of these applications, the boxes labeled “Batch Submit” (color-coded in red and green) themselves invoke substantial subworkflows. The two subworkflows leverage Globus Genomics features to launch multiple processes in parallel on the Amazon cloud, for (B) replicate alignment and (D) biosample footprint generation, respectively.

We mention a few additional features of the workflow. The first application in the master workflow, “Get BDBag from Minid,” dereferences a supplied Minid to retrieve the contents of the BDBag that it identifies. Thus the workflow can operate on any DNase-seq dataset contained in a BDBag and referenced by a Minid, including but not restricted to those produced by the **`encode2bag`** service.

This third application in the master workflow, “SNAP Workflow Batch,” invokes a subworkflow that comprises five applications (see Fig 5B). This subworkflow resolves the BDBag to identify the number of biosamples and replicates for an input tissue. Its second step manages a second subworkflow, Fig 5C, to execute the SNAP alignment algorithm for the input replicates of a biosample, All replicate alignments are executed in parallel and monitored for completeness. Once all replicate BAM files of a biosample are generated, the workflow merges and sorts them to produce a single BAM file for the biosample.

##### 4.3 Identifying footprints

Having assembled the DNase-seq data into a set of aligned BAM files, ❸ of Fig 1 uses the F-Seq program [34] to identify regions of open chromatin and then applies the HINT and Wellington footprint algorithms to those regions to generate footprints. This logic is implemented by the lower three applications in the master workflow shown in Fig 5A. The “Footprints Workflow Batch Submit” application runs the footprint generation subworkflow of Fig 5D, which first converts BAM files to Browser Extensible Data (BED) format, as required by the F-Seq tool; then runs the F-Seq tool on the BED file to identify areas of open chromatin; and finally runs the Wellington and HINT footprinting algorithms on both BED and BAM files to generate candidate footprints. Additional information on the generation process is available online [35].

##### 4.4 Generating the catalog of hits

While each footprint identified in ❸ is a *potential* TFBS, simply calling each footprint as a TFBS does not identify the cognate TF. Additionally, some footprints may correspond to unidentified or uncharacterized TFs that cannot be utilized. In preparation for eliminating such footprints, ❹ creates a *hit catalog* that links the observed footprints to known TF binding motifs.

The input to ❹, shown as Motif in Fig 1, is a collection of 1,530 nonredundant TF binding motifs assembled by Funk et al. [13]. This motif collection was assembled by using the Tomtom program from the MEME suite [36] to identify non-redundant motifs within the JASPAR 2016 [37], HOCOMOCO v10 [38], UniPROBE [39], and SwissRegulon [40] motif libraries, each of which was accessed via the Bioconductor R package MotifDb [41]. Tomtom was then used to compute pair-wise simularity scores for motifs from different libraries and then used those scores to eliminate redundant motifs. More details are available online [35]. This process involves human judgment and so we do not record the associated code as part of our reproducibility package. Rather we make available the resulting human-curated catalog to enable reproducibility of the subsequent workflow.

❹ uses the Find Individual Motif Occurrences (FIMO) tool [42], also from the MEME suite, to identity potential TF binding sites in the GRCh38 human reference genome. It uses FIMO (from the Regulatory Genomics toolkit version 0.11.2 as captured in the Docker container **`minid:b9jd6f`**) to search GRCh38 (captured in the hg38 folder of **`minid:b9fx1s`**) for matches with each of the 1,530 non-redundant motifs. An individual motif can match multiple times, and thus the output of this step is a total of 1,344,953,740 hits, each comprising a motif, a genomic location, the probability of the motif occurring at that location, and the match score of the sequence position.

##### 4.5 TFBS inference

The final step in the TFBS workflow involves intersecting the footprints produced in ❸ with the hits produced in ❹ to generate a set of candidate TFBSs. To accelerate the process of intersecting genomic locations, the Bioconductor R package GenomicRanges [43] is used to create GRanges objects for each of the 108 footprint files and for the hits catalog. Each footprint file is then intersected with the hits catalog independently to produce a total of 108 TFBS candidate files. For convenience, the footprints and TFBSs are also loaded into a cloud-based relational database, organized by tissue type, accessible as described online [44].

#### 5 Recap: A FAIR TFBS workflow

We review here the complete TFBS workflow, for which we specify the input datasets consumed by the workflow, the output datasets produced by the workflow, and the programs used to transform the inputs into the outputs. The inputs and programs are provided to enable readers to reproduce the results of the workflow; the outputs are provided for readers who want to use those results.

We specify each input, output, and program by providing a Minid. Several of these Minids reference what we call a “bag of bags” BDBag: a single BDBag that itself contains a set of BDBags, for example one per tissue. This use of a bag of bags allows us to refer to the dataset with a single identifier; the reader (or a program) can access the complete dataset by retrieving the bag of bags and using the BDBag tools to automatically download and materialize the constituent BDBags contained in its data directory. Each BDBag contains descriptive metadata for its contents.

Table 2 provide identifiers for the six datasets shown in Fig 1, and Table 3 provides identifiers for the software used to implement the five computational steps of Fig 1. We also aggregate the information in these tables into a single document so that they can be accessed via a persistent digital object identifier [45]. To simplify the creation of Docker container components, we created a tool that generates a Docker manifest from a Galaxy tool definition [46].

**Table 2.** The six datasets shown in Fig 1, D1–D6. For each we indicate whether it is an input or output.

| # | Name | Identifier | Role | Description | Size |
| --- | --- | --- | --- | --- | --- |
| D1 | DNase-seq | minid:b9dt2t | In | BDBag of 27 BDBags extracted from ENCODE by ❶, one per tissue: 1,591 FASTQ files in all. | 2.40 TB |
| D2 | Alignment | minid:b9vx04 | Out | BDBag of 54 BDBags produced by ❷, 1 per {tissue, seed}: 386 BAM files in all. | 5.30 TB |
| D3 | Footprints | minid:b9496p | Out | BDBag of 54 BDBags containing footprints computed by ❸, one per {tissue, seed}. Each BDBag contains two BED files per biosample, one per footprinting method. | 0.04 TB |
| D4 | Motifs | minid:b97957 | In | Database dump file containing the non-redundant motifs provided by Funk et al. [13]. | 31.5 GB |
| D5 | Hits | minid:b9p09p | Out | Database dump file containing the hits produced by ❹. | 0.04 TB |
| D6 | TFBSs | minid:b9v398 | Out | BDBag of 54 BDBags containing candidate TFBSs produced by ❺, one per {tissue, seed}. Each BDBag contains two database dump files, one per footprinting method. | 0.35 TB |

**Table 3.** The software used to implement the five steps shown in Fig 1. As the software for ❶ is used only to produce the input data at **`minid:b9dt2t`**, we do not provide identifiers for specific versions of that software.

| # | Name | Identifiers for software |
| --- | --- | --- |
| ❶ | Extract DNase-Seq | encode2bag service: https://github.com/ini-bdds/encode2bag-service<br>encode2bag client: https://github.com/ini-bdds/encode2bag |
| ❷, ❸ | Alignment, Footprints | Galaxy pipeline: minid:b93m4q<br>Dockerfile: minid:b9jd6f<br>Docker image: minid:b97x0j |
| ❹ | Hits | R script: minid:b9zh5t |
| ❺ | TFBSs | R scripts: minid:b9fx1s |

As a first step towards evaluating whether this information is enough to enable reproducibility, we asked a colleague to reproduce an analysis described in this paper: specifically, to regenerate the results for urinary bladder, for which, as shown in Table 1, there are only two replicates. They were successful.

#### 6 Discussion

The TFBS inference workflow implementation presented in Section 4 is structured in a way that it can be easily re-run by others. It is, furthermore, organized in a way that allows it to make easy use of parallel cloud computing. These desirable properties are the result of a disciplined approach to application development that aims for compliance with the ten simple rules for reproducible computational research defined by Sandve et al. [47]:

1. *For every result, keep track of how it was produced.* We preserve workflows and assign Minids to workflow results.
2. *Avoid manual data manipulation steps.* We encode all data manipulation steps in either Galaxy workflows or R scripts.
3. *Archive the exact versions of all external programs used.* We create a Docker container with versions of the tools used in the analysis, and generate Minids for the Docker file and Docker image of the container.
4. *Version control all custom scripts.* We maintain our programs in GitHub, which supports versioning, and provide Minids for the versions used.
5. *Record all intermediate results, when possible in standardized formats.* We record the major intermediate results, in the same manner as inputs and output, using FASTQ, BAM, and BED formats. In the case of database files, we dump tables to a text file via SQL commands.
6. *For analyses that include randomness, note underlying random seeds.* F-Seq uses the Java random number generator, but does not set or record a seed. We would need to modify F-Seq to record that information.
7. *Always store raw data behind plots.* Minids provide concise references to the raw data used to create the plots in the paper, which are bundled in BDBags.
8. *Generate hierarchical analysis output, allowing layers of increasing detail to be inspected.* Because we record the complete provenance of each result, a reader can easily trace lineage from a fact, plot, or summarized result back through the processing steps and intermediate and raw data used to derive that result.
9. *Connect textual statements to underlying results.* Our use of Minids would make it easy for Funk et al. [13] to reference specific data in their text. They do not do at present, but may in a future version of their paper.
10. *Provide public access to scripts, runs, and results.* Each is publicly available at a location accessible via a persistent identifier, as detailed in Tables 2 and 3.

The tools used in this case study do not in themselves ensure reproducibility and scalable execution, but they make it easy to create an implementation with those characteristics. For example, BDBag tools and Minid and Globus APIs allowed us to create the **`encode2bag`** web interface to ENCODE in just a few hours of effort, permitting a streamlining of the overall workflow that we likely would not have attempted otherwise. Similarly, the ability to create a new Minid at any time via a simple API call made it straightforward to create persistent identifiers for intermediate data products, which contributes to those data being Findable, Accessible, Interoperable, and Reusable (FAIR)—four attributes of digital objects that are often viewed as fundamental to data-driven discovery [5]. And the fact that we could easily create a readable specification of the ensemble footprinting method as a Galaxy workflow, and then dispatch that workflow to Globus Genomics for parallel cloud execution without regard to the location of input and output data, reduced both time requirements and opportunities for error in those logistically complex computations. So too did the ease with which we could package applications within Docker containers.

##### 6.1 Other approaches

It is instructive to compare and contrast the methods described in this paper with other approaches to big data and/or reproducible science.

Biomedicine is not alone in struggling with the complexities described here [48]. But big data tools from outside biomedicine tend to focus on narrow aspects of the analytic problem, leaving researchers on their own when it comes to managing the end-to-end discovery process [49].

Many approaches to reproducibility focus on using mechanisms such as makefiles [50,51], open source software [30,31], specialized programming environments [52], and virtual machines [53] to organize the code and/or data required for a computation. These approaches work well for small data but face challenges when computations must scale to terabytes and span sites.

Another set of approaches require that all data be placed, and analysis occur, within a single, homogeneous environment. In the case of the Analysis Commons [54] and Genomic Data Commons [55], this environment is a (public or private) cloud. Other systems leverage containers and related technologies to create a single reproducible artifact. Binder [56] allows researchers to create computational environments to interact with published code and data. Interactive notebooks, housed in public GitHub repositories, can be run in a version-controlled computational environment. Researchers structure their repository following simple conventions and include build files to describe the computational environment. Binder then uses a JupyterHub-like model to construct and spawn a computational environment in which to execute the notebook. Similarly, WholeTale [57] allows a researcher to construct a “Tale,” a computational narrative for a result. The researcher constructs a computational environment, selects one or more frontend analysis environments (e.g., Jupyter), and conducts their research within that environment. WholeTale tracks data imported into the environment (via linkage to identifiers or checksums) to produce a reproducible artifact (data, computation, and environment) for subsequent reuse and verification.

These approaches reduce complexity by enforcing physical or logical locality. They can work well when all required data and code can be integrated into the required homogenous environment. However, as the TFBS case study illustrates, data and computation are often distributed. Furthermore, the ability to move seamlessly among different storage and computational environments, as enabled by tools such as Globus, BDBags, and Globus Genomics, increases flexibility. The approach presented in this paper represents an alternative strategy for making science reproducible by directly addressing the needs of researchers working in loosely coupled environments in which multiple tools, services, and scripts are combined with distributed data products to conduct a given analysis.

##### 6.2 A data commons

Rather than requiring the use of a single computational environment, the technologies used in this case study facilitate interoperability among environments, so that data can be accessed from many locations (Globus Connect) using common security mechanisms (Globus Auth), transferred in a compact form (BDBags) with consistent naming and checksums for verification of integrity (Minids), and then analyzed rapidly using software in common formats (Docker), declarative workflows (Galaxy), and parallel computation (Globus Genomics). These elements represent useful steps towards a data commons, which Bonnazi et al. [58] have described in these terms:

> a shared virtual space where scientists can work with the digital objects of biomedical research; i.e., it is a system that will allow investigators to find, manage, share, use, and reuse data, software, metadata and workflows. It is a digital ecosystem that supports open science and leverages currently available computing platforms in a flexible and scalable manner to allow researchers to find and use computing environments, access public data sets and connect with other resources and tools (e.g. other data, software, workflows, etc.) associated with scholarly research.

By thus reducing barriers to finding and working with large data and complex software, our strategy makes it easier for researchers to access, analyze, and share data without regard to scale or location.

#### 7 Summary

We have presented tools designed to facilitate the implementation of complex, “big data” computations in ways that make the associated data and code findable, accessible, interoperable, and reusable (FAIR). To illustrate the use of these tools, we have described the implementation of a multi-stage DNase I hypersensitive sites sequencing data analysis that retrieves large datasets from a public repository and uses a mix of parallel cloud and workstation computation to identify candidate transcription factor binding sites. This pipeline can be rerun in its current form, for example as new DNase I hypersensitive sites sequencing data become available; extended with additional footprinting methods (for example, protein interaction quantification [59]) as new techniques become available; or modified to apply different integration and analysis methods. The case study thus demonstrates solutions to problems of scale and reproducibility in the heterogeneous, distributed world that characterizes much of modern biomedicine. We hope to see others experiment with these tools in other contexts and report their experiences.

## Acknowledgments

We thank the Galaxy and Globus teams, and other developers of tools on which we build here, for their work. We also thank the NIH Commons Cloud Credits program and the Amazon Web Services Research Cloud Credits program for their support. This work was supported in part by NIH contracts 1U54EB020406-01: Big Data for Discovery Science Center, 1OT3OD025458-01: A Commons Platform for Promoting Continuous Fairness, and 5R01HG009018: Hardening Globus Genomics; and DOE contract DE-AC02-06CH11357.

